# Estimation of Average Blood Glucose Values Based on Fructosamine Values

**DOI:** 10.1101/2021.07.25.453711

**Authors:** Luis Jesuino de Oliveira Andrade, Alcina Maria Vinhaes Bittencourt, Luiz Felipe Moreno de Brito, Luís Matos de Oliveira, Gabriela Correia Matos de Oliveira

## Abstract

**Introduction:** The fructosamine is originated of the glycation of plasmatic proteins, especially albumin, in addition to immunoglobulins and proteins diverse. It constitutes an alternative biomarker of glycemic control when glycated hemoglobin is not indicated for this purpose.

**Objective:** To define the mathematical relationship between fructosamine and average glucose values.

**Method:** The study comprised the laboratorial data collected of 1227 diabetic subjects (type 1 and type 2). Fructosamine levels obtained at the end of three weeks and measured were compared with the average glucose levels of the three previous weeks. The average glucose levels were determined by the weighted mean of the daily fasting capillary glucose results performed during the study period, and the plasma glucose taken at the time of the fructosamine.

**Results:** A total of 9,450 glucoses were performed. Linear regression analysis between the fructosamine and average glucose levels showed that each increase of 1.0 µmol/L in fructosamine increase 0.5mg/dL in the average glucose levels as evidenced in the equation forward: Average glucose levels = 0.5157 x Fructosamine - 20. According to the coefficient of determination (r2 = 0.353492, P < 0.006881), making it possible to calculate the estimated average glucose according to the frutosamine values.

**Conclusion:** Fructosamine levels can be expressed as average glucose levels for assessing the metabolic control of diabetic patients.

## INTRODUCTION

Fructosamine (1-amino-1-deoxy fructose) is an expression used for all glycated plasmatic proteins, especially albumin which represents 55-80% of the glycosylated proteins in serum, in addition to immunoglobulins and proteins diverse. This is a ketoamine constituted through an irreversible nonenzymatic connection of glucose with plasmatic proteins via glycation. The glycation process corresponds to the initial formation of a Schiff base (aldimine) which is subsequently rearranged into a stable Amadori product (ketoamine) as a function of the binding of the amino acids cysteine, arginine, and lysine to glucose within the Schiff base (aldimine) rearranged later into a stable product Amadori (ketoamine) as a function of the binding of the amino acids cysteine, arginine and lysine to glucose within the protein molecules.^1^

The fructosamin constitutes an alternative biomarker of glycemic control when glycated hemoglobin is not indicated for this purpose, and evaluates glucose control over a 2–3 week period.^2^

The fact that fructosamine to reflects the average glucose over a 2-3 week period allows for more accurate glycemic control and better therapeutic adjustment, especially in unstable diabetes mellitus (DM), therefore, more suitable for monitoring the response to treatment.

Fructosamine has not been widely used as compared to glycosylated hemoglobin (HbA1c) for monitoring the metabolic control of DM, although several reports assure that fructosamine could be superior to HbA1c.^3^ In addition, fructosamine is unaltered by hemoglobinopathies (structural hemoglobin variants and thalassemia syndromes) or anemia as in case with HbA1c, and thus can be employed in circumstances where HbA1c is unsafe to evaluate due to biological or analytical interferences.^4^ A study comparing fructosamine and HbA1c in diabetic subjects and normal controls showed a good correlation between the two parameters in diabetic subjects, but not in normal subjects.^5^

Some time ago, the clinical use of fructosamine was validated as a standard of short-term glycemic control in diabetic patients.^6^ Thus the objective of this study was to define the mathematical relationship between fructosamine and average glucose values.

## METHODS

The study comprised the laboratorial data collected of 1227 diabetic subjects (DM type 1 and DM type 2). Fructosamine levels measured and obtained at the end of three weeks were compared with the average glucose levels of the three previous weeks. The average glucose levels were determined by the weighted mean of the daily fasting capillary glucose results performed during the study period, and the plasma glucose taken at the time of the fructosamine.

Statistical analysis was carried out by Excel and in order to investigate meaningful correlation between serum fructosamine and average glucose levels. We use linear regression to evaluate the mathematical relationship between fructosamine levels and average glucose levels. In all calculations p<0.05 was considered as statistically significant.

## RESULTS

A total of 9,450 glucoses and 1,227 fructosamine were performed. Linear regression analysis between the fructosamine and average glucose levels (Table 1) showed that each increase of 1.0 µmol/L in fructosamine increase 0.5mg/dL in the average glucose levels as evidenced in the equation forward: Average glucose levels = 0.5157 x Fructosamine - 20.

**Table 1.** * Average glucose levels = 0.5157 x Fructosamine - 20.

| Fructosamine<br>$\mu\text{mol/L}$ | Estimated average glucose<br>mg/dl* |
| --- | --- |
| 205 | 80.0 |
| 215 | 90.8 |
| 225 | 96.0 |
| 235 | 101.1 |
| 245 | 106.3 |
| 255 | 111.5 |
| 265 | 116.6 |
| 275 | 121.8 |
| 285 | 126.9 |

According to the coefficient of determination (r2 = 0.353492, P < 0.006881) (Table 2), making it possible to calculate the estimated average glucose according to the frutosamine values (Table 3).

**Table 2.** Coefficient of determination.

| *Regression Statistics* |  |
| --- | --- |
| Multiple R | 0,594552 |
| R-square | **0,353492** |
| Adjusted R-square | 0,352045 |
| Standard error | 49,30802 |
| Observations | 1227 |

**Table 3.**
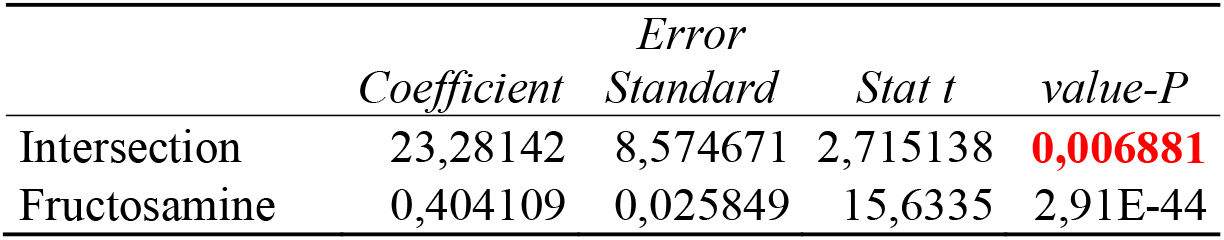

The relationship between the average glucose and fructosamine is present in figure 1.

**Figure 1.**
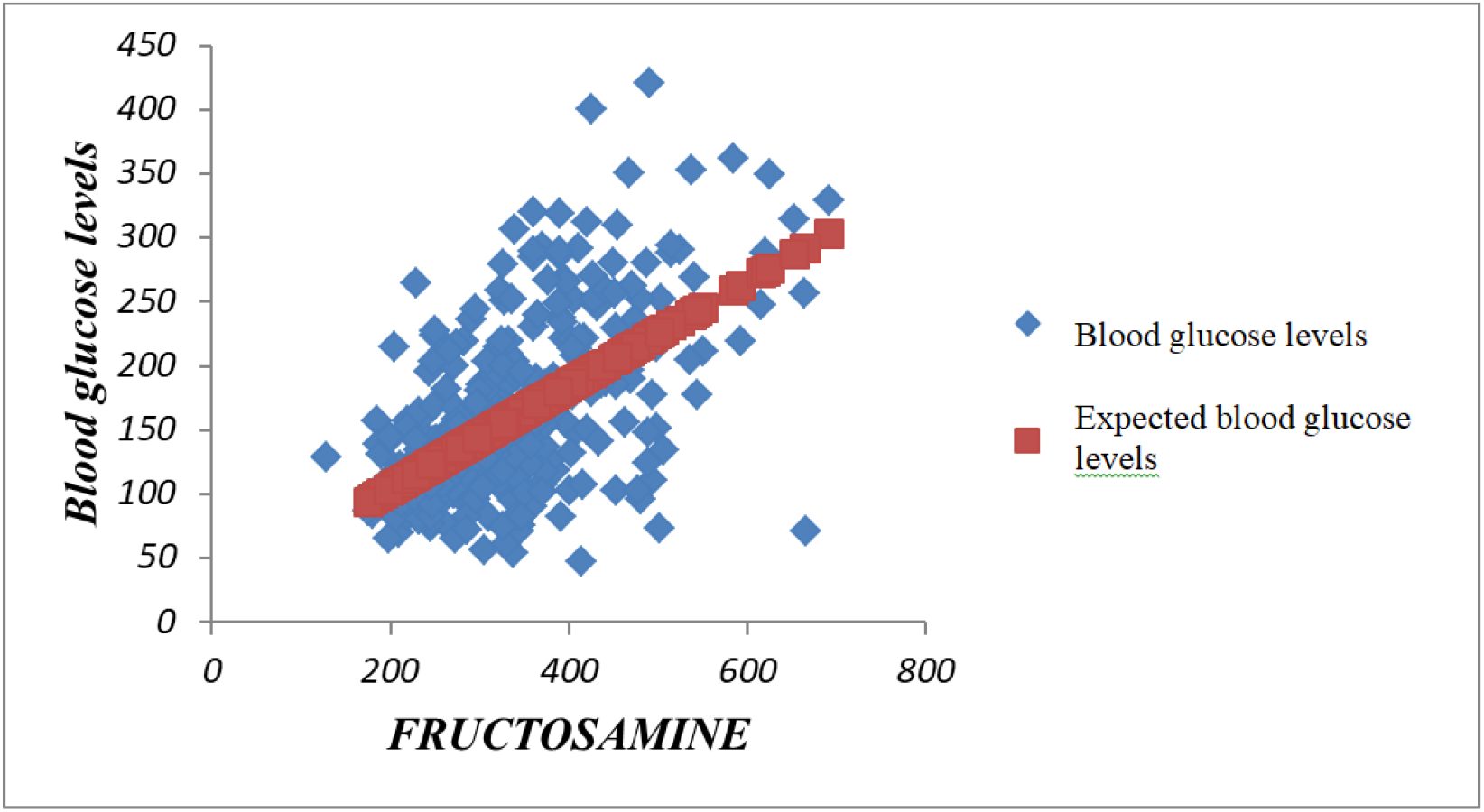
Relationship between the baseline average glucose and fructosamine.

## DISCUSSION

In this study, we define the mathematical relationship between fructosamine and average glucose values established by the following equation: Average glucose levels = 0.5157 x Fructosamine - 20.

Human albumin is extracellular multifunctional protein, and became more recently a biomarker of hyperglycemia. Fructosamine is a reliable biomarker of glycemic control, having a low cost and easy laboratory evaluation, showing a good correlation with the average plasma glucose over a period of 2-3 weeks. Thus, the fructosamine is a term attributed to all ketoamine linkages resulting in glycation of serum proteins.^7^ Unlike frutosamine, which provides short-term (2-3 weeks prior) glycemic control information, HbA1c provides long-term (2-3 months prior) average glycemic control information.^8^ The rate of non-enzymatic glycation of frutosamine, however, is about 9 to 10 times higher than that of Hba1c.

The criteria for the diagnosis of DM according to the American Diabetes Association include: HbA1C criteria or plasma glucose criteria (fasting plasma glucose or the 2-h plasma glucose value after a 75-g oral glucose tolerance test).^9^

A study to define the mathematical relationship between Hb1c and the average blood glucose over the previous three months in diabetics defined an equation to calculate the average blood glucose, where the average blood glucose = 28.3 × A1C −43.9.^10^ Our study using linear regression analysis, correlating frutosamine levels and mean glucose levels, observed that for each increase of 1.0 µmol/L there was a 0.5 mg/dL increase in mean glucose levels, as evidenced in the forward equation: Mean glucose levels = 0.5157 x Fructosamine - 20. Thus, according to the coefficient of determination it becomes possible to calculate the estimated mean glucose according to the fructosamine values. Our results give support the linear relationship between fructosamina and avarage glucose.

There is growing interest in the use of frutosamine in screening and follow-up of DM metabolic control. The evaluation of fructosamine has been proposed to upgrade diagnosis and to monitor of DM, as a function of the pitfalls that can occur with HbA1c.^11^ Additionally, the assessment of fructosamine has been proposed as a predictor of DM risk.^12^ Thus, fructosamine is closely associated with diabetes risk and its elevation can be a useful indicator of a future risk of DM, independent of glucose measurements.

A normal value of fructosamine varies of 200-285 µmol/L, when the serum albumin concentration level is 5 g/dL, and the manufacturer’s reference value interval varies of 205–285□µmol/L.^13^ A study to estimate the prevalence of DM from frutosamine assessment associated with the glucose tolerance test with 76 grams of glucose, using a frutosamine of 310 µmol/L as the cut-off point for DM diagnosis, showed that the standard error rates of estimated prevalence of DM ranged from 40% for a population of 200 individuals to 20% for a population of 2,000 individuals or more.^14^ Our results showed that the equation obtained allows us to calculate the average blood glucose to frutosamine values in diabetic individuals, thus assessing their short-term glycemic control.

## CONCLUSION

Fructosamine levels can be expressed as average glucose levels for assessing the metabolic control of diabetic patients. However, further studies are needed to validate the relationship of frutosamine levels and average blood glucose levels.

